# CLM-access: A Specialized Foundation Model for High-dimensional Single-cell ATAC-seq analysis

**DOI:** 10.1101/2025.08.10.669570

**Authors:** Ziqiang Liu, Bowen Li, Zhenyu Xu, Yantao Li, Junwei Zhang, Chulin Sha, Xiaolin Li

## Abstract

Inspired by the success of large language models (LLMs) in natural language processing, cell language models (CLMs) have emerged as a promising paradigm for learning cell representations from high-dimensional single-cell data—particularly transcriptomic profiles from scRNA-seq. These foundation models have shown remarkable potential across a variety of downstream applications. However, there remains a lack of foundation models for scATAC-seq data, which measures chromatin accessibility at single-cell level and is critical for decoding epigenetic regulation. Developing such models is considerably more challenging due to the unique characteristics of scATAC-seq data, including the vast number of chromatin regions, lack of standardized annotations, extreme sparsity, and near-binary distributions. To address these challenges, we systematically explore various strategies and propose CLM-access, a specialized foundation model for scATAC-seq data. CLM-access incorporates three main innovations: (1) an unified data processing pipeline that maps 2.8 million cells onto an unified reference of over 1 million chromatin regions; (2) a specialized patching and embedding strategy to effectively manage high-dimensional inputs; and (3) a tailored masking and loss function design that preserves fine-grained regional information while enhancing training efficiency and representation quality. With comprehensive benchmarks, we show that CLM-access significantly outperforms existing methods in key downstream tasks, including batch effect correction, cell type annotation, RNA expression prediction, and multi-modal integration. This work establishes a scalable and interpretable foundation model for single-cell epigenomic analysis and expands the application of CLMs in single-cell research. Code is available at https://github.com/HIM-AIM/CLM-access

## 1 Introduction

The rapid advancement of single-cell omics technologies has resulted in an explosion of high-dimensional data, posing challenges in effectively delineating meaningful biological insights. The scATAC-seq (single-cell Assay for Transposase-Accessible Chromatin using sequencing) data analysis is a typical example. This technique is used to assess open chromatin regions of a cell[1], which enables the localization of active cis-regulatory elements (CREs) such as promoters, enhancers, etc., thereby elucidating the dynamics of gene regulation in various cell types[2, 3, 4]. In recent years, over a few large-scale scATAC-seq data atlases have been generated[5, 6], offering resources for investigating CREs and their regulating genes during key biological processes such as embryonic development and brain regional differentiation [7, 8, 9]. However, scATAC-seq data analysis encounters several primary challenges: First, the scale of candidate CREs (cCREs) reaches 10^5^-10^6^ orders of magnitude, resulting in a drastic increase in information encoding dimension; Second, the data exhibit extreme sparsity, leading to a discrete distribution of signals across the genome; Third, the binary nature of the signals (accessible/inaccessible) makes it difficult to directly reflect the interaction hierarchies of the CREs[10, 11, 12]. In addition, unlike scRNA-seq data where standard gene reference is available, scATAC-seq data lacks universally defined genome locations or annotations of cCREs, which makes integrating scATAC-seq data from different experiments particularly challenging[13, 14]. These characteristics collectively constrain the scATAC-seq data analysing methods development in revealing gene regulatory networks[15, 16, 17].

Current scATAC-seq data analytical tools can be categorized into task-specific methods and integrated analytical pipelines[18, 19]. Task-specific methods exhibit limited generalizability, requiring substantial reconfiguration of model architectures to adapt to new scenarios[20, 21]. Whereas integrated methods despite incorporating multi-module functionalities, often involve a high proportion (typically exceeding 80%) of pre-filtering of cCREs[22, 23]. This result in dimensionality reduction and information entropy loss in the data, thereby undermining the model’s capacity to decipher complex regulatory relationships[24, 25].

More recently, several cell language foundation models have been successfully developed for scRNA-seq data, including scBERT[26], Geneformer[27], scGPT[28], and scFoundation[29]. These approaches treat genes as tokens and transform scRNA-seq data into a language-like format. Leveraging large-scale datasets and self-supervised pretraining strategies, these models can learn generalizable representations of cells that facilitate data integration and biological interpretation. They also exhibit strong potential in downstream tasks such as cell type annotation, biomarker discovery, and in silico gene perturbation analysis, etc. These studies provide encouraging evidence that a foundation model tailored for scATAC-seq data could address key limitations of existing methods and offer a more powerful framework for interpreting epigenetic regulation at single-cell resolution. However, there is still much to explore in terms of how to best represent, train, and scale models that are well-suited for the complexity of scATAC-seq data.

In response to these unmet needs, we present CLM-access—a Transformer-based cell language foundation model specifically designed for scATAC-seq data. We begin by establishing a standardized data processing pipeline to construct an integrated Human-scATAC-seq dataset encompassing approximately 2.8 million cells for model pretraining. To address the challenges inherent in scATAC-seq data, we systematically explored multiple strategies for tokenization, masking, and loss function design. To handle the high dimensionality, we partitioned accessible chromatin regions into patches, each consisting of a fixed number of peaks (corresponding to cCREs mapped in the genome), and treated each patch as a token. The model inputs combine token embeddings with peak-level representations and are processed through a Transformer architecture to perform masked peak reconstruction, optimized using binary cross-entropy (BCE) loss. This design allows the model to incorporate all cCREs as input, effectively accommodate data sparsity, and simultaneously ensure high training efficiency and robust learning performance. The final CLM-access architecture comprising roughly 20 million parameters, includes an embedding module, a Transformer encoder, and a peak decoder. With fine-tuning, CLM-access achieves considerably better performance than existing methods on key tasks including batch effect correction, cell type annotation, RNA expression prediction and multi-modal integration after fine-tuning, providing a powerful tool for scATAC-seq data research.

## 2 Related Work

### 2.1 Deep-learning Methods for Analyzing scATAC-seq Data

The majority of deep learning methods for scATAC-seq data analysis are based on variational autoencoder (VAE) or Transformer architectures, and often rely on paired scRNA-seq and scATAC-seq datasets for multi-modal integration. Methods such as scButterfly and MultiVI adopt VAE frameworks to perform multi-omics analysis and integration[30, 31], but they pre-filter the input by retaining only highly variable peaks to lower the data dimensionality. More recently, scCLIP has emerged as an innovative Transformer-based model for integrating scRNA-seq and scATAC-seq data, and partitions peaks into patch regions for feature extraction. However, it is trained on a relatively small data[32].

### 2.2 Foundation Models for scRNA-seq Data

Geneformer, scGPT, and scBERT are foundational models pre-trained on millions of single-cell RNA sequencing (scRNA-seq) profiles, demonstrating exceptional performance in tasks such as cell type annotation and gene network inference [27, 28, 26]. Both Geneformer and scBERT draw inspiration from the BERT model architecture, of which Geneformer innovatively incorporates gene ordering processing. In contrast, models like scGPT, utilize a generative pre-training approach for training. All these models treat cells as “gene sentences” to predict the expression of hidden genes. However, the direct transfer of models trained on scRNA-seq data to single-cell chromatin accessibility sequencing (scATAC-seq) data presents challenges due to the vast number of cCREs, extreme sparsity, and ambiguous regulatory relationships inherent in scATAC-seq data.

## 3 Methodology

### 3.1 Overview of CLM-access

As depicted in Figure 1, this study employs the BERT paradigm for single-cell assay for transposase-accessible chromatin using sequencing (scATAC-seq) analysis. The model leverages self-supervised pre-training and the core Transformer architecture to simulate long-range chromatin interactions. In terms of embedding processing, the model optimizes the token design in BERT to accommodate the peak characteristics of scATAC-seq. Specifically, it partitions high-dimensional ATAC-seq data (which can reach millions of dimensions) into 2,000 fixed-size patches, with each segment encoded as a token. This process compresses the peak information within each segment into a low-dimensional representation, thereby creating specialized token embeddings for chromatin accessibility data. This approach addresses the issue encountered by traditional ATAC-seq models, where the extreme sparsity of the data renders it infeasible to input all data at once. It preserves the complete information of the data while enabling adaptation to the Transformer architecture. Traditional methods, such as performing TF-IDF transformation or removing zero values, inevitably result in information loss. Particularly, TF-IDF transformation confines the data to the training set, necessitating recalculation when additional training data is incorporated, thereby compromising the model’s scalability in subsequent applications.

**Figure 1.**
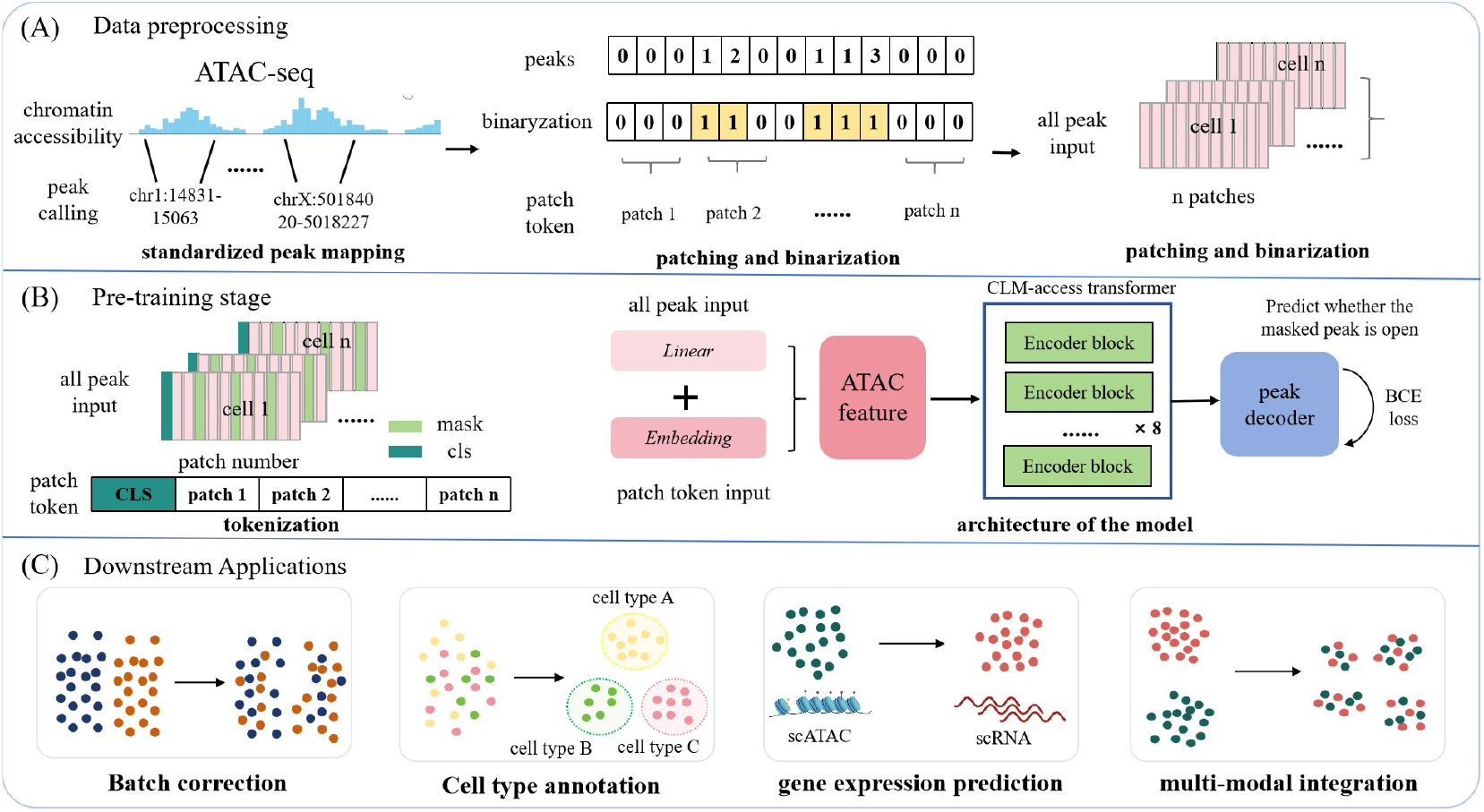
Overview of CLM-access. **A**, The tokenization process in CLM-access involves processing the accessible chromatin open regions from scATAC-seq data and dividing them into multiple regions to form cellular sentences. **B**, The model architecture of CLM-access consists of three modules: (1) an embedding module that converts cell sentences and peak matrices into embedding sequences through embedding layers and linear layers, respectively, and then sums them up; (2) the CLM-access transformer, which includes multiple transformer blocks utilizing a attention mechanism to generate cell embeddings and contextual region embeddings; (3) a peak decoder that reconstructs whether a peak is accessible from the cell embeddings. **C**,The downstream tasks of CLM-access include batch effect removal, cell type annotation, gene expression prediction, and multi-modal integration.

The global receptive field of the Transformer enables it to learn patterns without relying on absolute positional data, thereby enhancing its performance on non-sequential patterns like those in scATAC-seq. By effectively adapting the BERT framework and incorporating specific optimizations in feature embedding and dimensionality reduction, this architecture provides a superior solution for scATAC-seq data.

### 3.2 Data Processing and Tokenization

We constructed the Human-scATAC database using 2.8M scATAC-seq data entries, preprocessed into a sparse cell-by-cCRE count matrix *X* ∈ *R* ^*N×P*^, where *N* and *P* denote the number of cells and the number of cCREs, respectively. More details on how the database is constructed is provided in the supplementary file. Directly modeling all 1.15M cCREs as tokens would inflate parameters and hinder training. To resolve this, we partitioned each cell into n patches, treating each patch as a language model token to aggregate regional cCRE signals. Additionally, we introduced a [cls] token at the cell’s start to capture its global chromatin accessibility context. The model thus processes each cell as a fixed-length patch sequence, structured as:

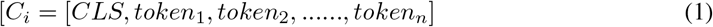

In the preprocessing step prior to patch division, all cCREs are systematically ordered based on their chromosomal positions. This sorting ensures a consistent and biologically meaningful arrangement of the cCREs, facilitating subsequent analysis and modeling steps. After determining the patch regions, we proceed to input all corresponding peak values within each patch into the model. To ensure uniformity across patches, we standardize the length of the peak values within each patch. Specifically, if the number of peak values in a patch is less than the predetermined uniform length, we pad the patch with additional values to meet this length requirement. The structure of each patch, including the handling of peak values and padding, is formally described by the following formula:

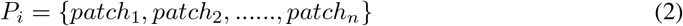

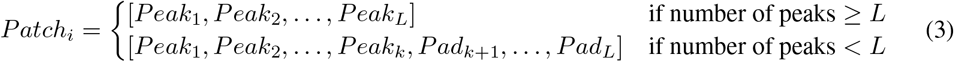

Here, *patch*_*i*_ represents the *i*-th patch, L is the predetermined uniform length for all patches, *Peak*_*j*_ denotes the *j*-th peak value within the patch, and *Pad*_*j*_ represents the padding value used to fill the patch to the desired length when the number of peak values is less than *L*. This approach ensures that each patch input to the model has a consistent structure, facilitating efficient processing and analysis. Finally, considering that the majority of peak values are either 0 or 1, with rare occurrences of values exceeding 2, a binarization process is applied to all peak values to facilitate model training. This step is taken to simplify the distribution of peak values, making it easier for the model to learn and generalize from the data. By converting the peak values into a binary form, we aim to enhance the training efficiency and effectiveness of the model.

### 3.3 Model Architecture of CLM-access

This subsection primarily delineates the pre-training architecture of the model, wherein downstream tasks are adapted through minor modifications and fine-tuning on the pre-trained framework. For detailed information, please refer to the supplementary materials.

#### 3.3.1 Peak Embedding Module

The embedding module comprises two primary components: the patch token embedding layer and the peak value embedding layer. The patch token embedding layer takes as input the processed sequence of patch tokens corresponding to each cell. These tokens represent distinct features or segments within the cell data. On the other hand, the peak value embedding layer receives the binarized and padded peak values as input. These peak values are organized into a matrix with a number of columns equal to the length of the cCRE index sequence for each cell, ensuring consistency in dimensionality across different inputs. Subsequently, the embedding module combines the outputs from these two embedding layers through element-wise addition. This integration can be formally represented as follows:

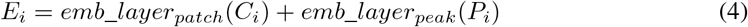

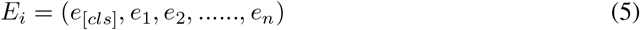

In the model, *emb*_*layer*1(·) and *emb*_*layer*2(·)) are the cCRE and ranking embedding layers, respectively. The patch token embedding layer mirrors NLP’s word embedding approach, but introduces a peak value embedding layer to enhance ATAC-seq data utilization. The embedding module uses a 256-dimensional space: the patch token layer encodes 1998 cCRE markers and 4 special tokens, while the peak value layer employs a linear transformation to align cellular matrix *P*_*i*_ with the patch token embedding dimension.

#### 3.3.2 Peak Encoder Module

The CLM-access Transformer, the core module of the Transformer-based foundational model, comprises 8 stacked blocks. These blocks employ a bidirectional attention mechanism to model global dependencies in high-dimensional data, capturing epigenetic regulatory networks across cells. Each block has an embedding dimension of 256, uses 8 attention heads, and processes both inputs and outputs, with their mathematical formulation as follows:

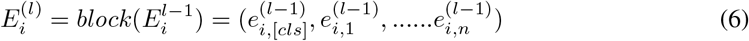

Here, 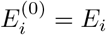, and 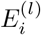 is the output of the *l*-th Transformer block, with *block*(·) as a single block. The Transformer captures patch interactions via bidirectional attention, described as:

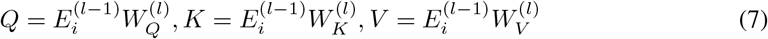

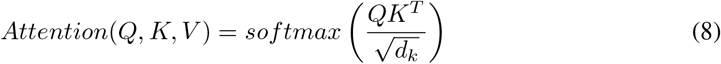

Wherein, 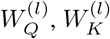, and 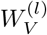 are trainable weight matrices utilized to compute the Query (Q), Key (K), and Value (V) respectively, with *d*_*k*_ denoting the dimension of the Key vector. To enhance the representational capability of the Transformer, each Transformer block employs a multi-head attention mechanism by computing multiple attention heads in parallel. Each attention head possesses its own set of 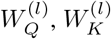, and 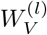, adhering to the same formulation as single-head attention. The outputs from all attention heads are concatenated and subsequently transformed linearly using a learnable matrix 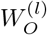, yielding the output embeddings. The output of the final Transformer block is denoted as 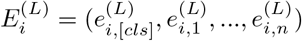, where 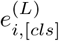 represents the cell embedding for cell *i*. The CLM-access Transformer comprises approximately 12 million parameters and is implemented using the Flash Attention v2 framework to ensure efficient training and inference.

#### 3.3.3 Peak Decoder Module

The peak decoder utilizes a single-layer, fully connected neural network to transform the cell embeddings into the values of all masked peaks for each cell. This process can be formulated as follows:

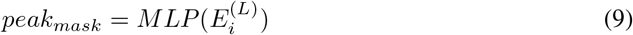

Finally, the predicted masked peak values and the true values are input into a Binary Cross-Entropy (BCE) loss function to train the model.

## 4 Experiment

This section explores the impact of various training strategies on pre-training performance. During the research, it was discovered that directly employing the training approaches of traditional scRNA-seq foundation models and conventional scATAC-seq deep learning models for training scATAC-seq foundation models is infeasible. Neither retaining only accessible peaks nor filtering for highly variable peaks as input can successfully train the model.

To this end, we dedicated substantial efforts to investigating training strategies and data input designs, leading to a series of meaningful insights that may serve as a theoretical foundation for the future development of foundational models for scATAC-seq data. Building upon the successful pre-training of CLM-access, we evaluated its performance across a range of downstream tasks, including batch effect mitigation, cell type annotation, RNA expression prediction, and cross-modality integration.

### 4.1 Experiment Settings

In our experimental design, we employed models with varying parameter scales and utilized datasets of distinct sizes. For the pre-training phase, all models were uniformly trained over 15 epochs with a batch size of 8. In contrast, the fine-tuning configurations for downstream tasks were tailored to each task’s specific requirements, leading to task-dependent adjustments. A comprehensive and detailed account of these experimental settings and their rationales can be found in the supplementary materials.

### 4.2 Comparison and Analysis of CLM-access Pre-training Module

This section primarily focuses on the critical pre-training factors of the CLM-access model, including whether to apply binarization, the selection of loss functions, and the forms of masking. We systematically investigated their impacts on the pre-training performance. In the experiment, we uniformly utilized a dataset of 370,000 single cells processed through the scCLIP pipeline [32]. This dataset was divided into a training set and a test set. We then calculated the Normalized Mutual Information (NMI) and Adjusted Rand Index (ARI) using the cell types in the test set for evaluation purposes.

#### 4.2.1 Binarization Processing and Patch Number Effects

We evaluated the performance of the data under different patch sizes and with the presence or absence of Topologically Associating Domains (TADs), comparing cases where the data was binarized versus when it was not. The experimental results are presented in Table 1. When only summing the intensity values of all peak signals without applying binarization processing, the model exhibited significant convergence difficulties during training and struggled to effectively capture the internal feature information of each patch region. In contrast, the introduction of binarization processing markedly reduced the model’s learning complexity. Further analysis revealed that as the granularity of patch regions decreased, the model retained more complete original information during the feature extraction process and demonstrated superior performance.

**Table 1:** The influence of data processing and patch size choice on the pre-trained model.

| metrics | processing | patch 1k | patch 2k | patch 5k | TAD(14k) |
| --- | --- | --- | --- | --- | --- |
| ARI | sum | 0.0007 | 0.0033 | 0.0001 | 0.0030 |
|  | binarization | 0.0113 | 0.0459 | 0.1688 | **0.2397** |
| NMI | sum | 0.0139 | 0.0212 | 0.0087 | 0.0206 |
|  | binarization | 0.0513 | 0.1617 | 0.4781 | **0.5752** |

#### 4.2.2 Loss Function Comparisons

In this study, we designed and compared two data processing strategies for plaque segmentation. In Strategy 1, peak signal intensities within each patch are summed then binarized, and the model is optimized using MSE and BCE loss functions, respectively, at the patch level. In Strategy 2, individual peak signals are binarized directly and input into the model on a patch-by-patch basis, with BCE loss applied at the peak level. We conducted a systematic comparison of the performance of these two approaches.

Initial results showed that reducing plaque granularity improved model performance, but summing signals within patches led to significant information loss. Increasing patch numbers mitigated this but raised computational costs due to longer input sequences. Hence, we adopted Strategy 2, binarizing all peak signals and processing them patch-by-patch.As Table 2 indicates, Strategy 2 achieved optimal performance with an input sequence length of only 2000. This approach not only enhanced model performance but also reduced training and inference time, effectively balancing computational efficiency and feature preservation.

**Table 2:** The influence of loss function choice on the pre-trained model.

| metrics | loss function | patch 1k | patch 2k | patch 5k |
| --- | --- | --- | --- | --- |
| ARI | patch-level MSE | 0.0113 | 0.0459 | 0.1688 |
|  | patch-level BCE | 0.0122 | 0.0394 | 0.0854 |
|  | peak-level BCE | 0.2120 | **0.2511** | 0.1304 |
| NMI | patch-level MSE | 0.0513 | 0.1617 | 0.4781 |
|  | patch-level BCE | 0.0537 | 0.1479 | 0.2560 |
|  | peak-level BCE | 0.5078 | **0.5813** | 0.3648 |

#### 4.2.3 Impact of Forms of Masking

Through systematic experimental design, this study explored the impact of different masking strategies on model performance, proposing two approaches: jointly masking patch tokens and peak signals, or masking only peak signals. We conducted comparative experiments to assess their differential effects on the model’s feature learning capabilities.

As shown in Table 3, the joint masking strategy substantially increased model complexity, hindering effective extraction of key biological features from highly masked input sequences. Consequently, subsequent experiments adopted the peak signal-only masking approach. This method maintained model convergence stability, reduced information loss, and enabled the model to focus on learning local features and global distribution patterns of peak signals. This optimization strategy provided crucial support for enhancing model performance in subsequent experiments.

**Table 3:** The influence of mask forms on the pre-trained model.

| metrics | mask forms | patch 1k | patch 2k | patch 5k |
| --- | --- | --- | --- | --- |
| ARI | mask token and value | 0.0019 | 0.0016 | 0.0048 |
|  | only mask value | 0.2120 | **0.2511** | 0.1304 |
| NMI | mask token and value | 0.0181 | 0.0181 | 0.0294 |
|  | only mask value | 0.5078 | **0.5813** | 0.3648 |

During pre-training, we evaluated the model’s zero-shot learning capability by varying parameter and dataset sizes. As Table 4 illustrates, models with moderate parameter counts exhibited optimal performance. Increasing parameters further yielded diminishing returns, suggesting a potential link to the pre-training dataset’s size and distribution. With limited data, excessively large models risk overfitting, thereby limiting generalization. Hence, we selected a medium parameter count for the Transformer component, balancing computational efficiency and model capability. We also conducted pre-training experiments with datasets of varying sizes. Table 4 shows that, with a consistent model architecture, increasing pre-training data volume led to a clear improvement in downstream task performance, approximately linearly correlated with data size. This finding underscores the enhancing effect of large-scale data on deep learning models’ feature representation capability, providing a theoretical foundation for dataset construction strategies in subsequent experiments.

**Table 4:**
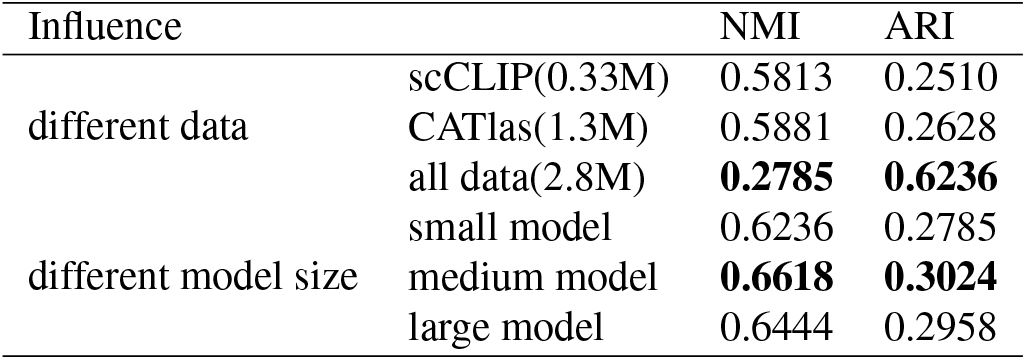
Model performances under different sizes of datasets and model parameters.

| Influence |  | NMI | ARI |
| --- | --- | --- | --- |
| different data | scCLIP(0.33M) | 0.5813 | 0.2510 |
|  | CATlas(1.3M) | 0.5881 | 0.2628 |
|  | all data(2.8M) | **0.2785** | **0.6236** |
| different model size | small model | 0.6236 | 0.2785 |
|  | medium model | **0.6618** | **0.3024** |
|  | large model | 0.6444 | 0.2958 |

### 4.3 CLM-access Excels in Key Downstream Tasks of scATAC-seq Analysis

In a typical downstream analysis pipeline for scATAC-seq data, key steps include batch effect removal, cell type annotation, and prediction of gene expression levels. These analytical tasks heavily rely on an accurate and robust computational analysis framework. CLM-access is a foundational model specifically tailored for scATAC-seq data, boasting high adaptability. As illustrated in Figure 1c, it can flexibly address diverse downstream analysis needs through fine-tuning and seamless integration with task-specific modules.

#### 4.3.1 Experimental Results of Batch Correction

Batch effects stem from systematic technical biases introduced during experimental processing, which may obscure biological differences among single cells and compromise the accuracy of downstream analyses. The pre-training data encompass samples from multiple batches, allowing the model to implicitly integrate batch relationships among cells. In this study, we take 4 PBMC scATAC-seq datasets (details in supplementary file) to validate the performance of batch effect removal, and compare CLM-access against two conventional methods PCA and harmony. As illustrated in Figure 2, in the zero-shot scenario, the CLM-access method outperforms traditional approaches such as PCA and Harmony [33]. After fine-tuning with batch information incorporated, the model’s performance is further significantly enhanced, surpassing its zero-shot performance. CLM-access is capable of constructing cell representations that are biologically meaningful and free from batch-related biases, all without relying on prior knowledge of cell types. This demonstrates its robust independence and broad applicability across various scenarios.

**Figure 2.**
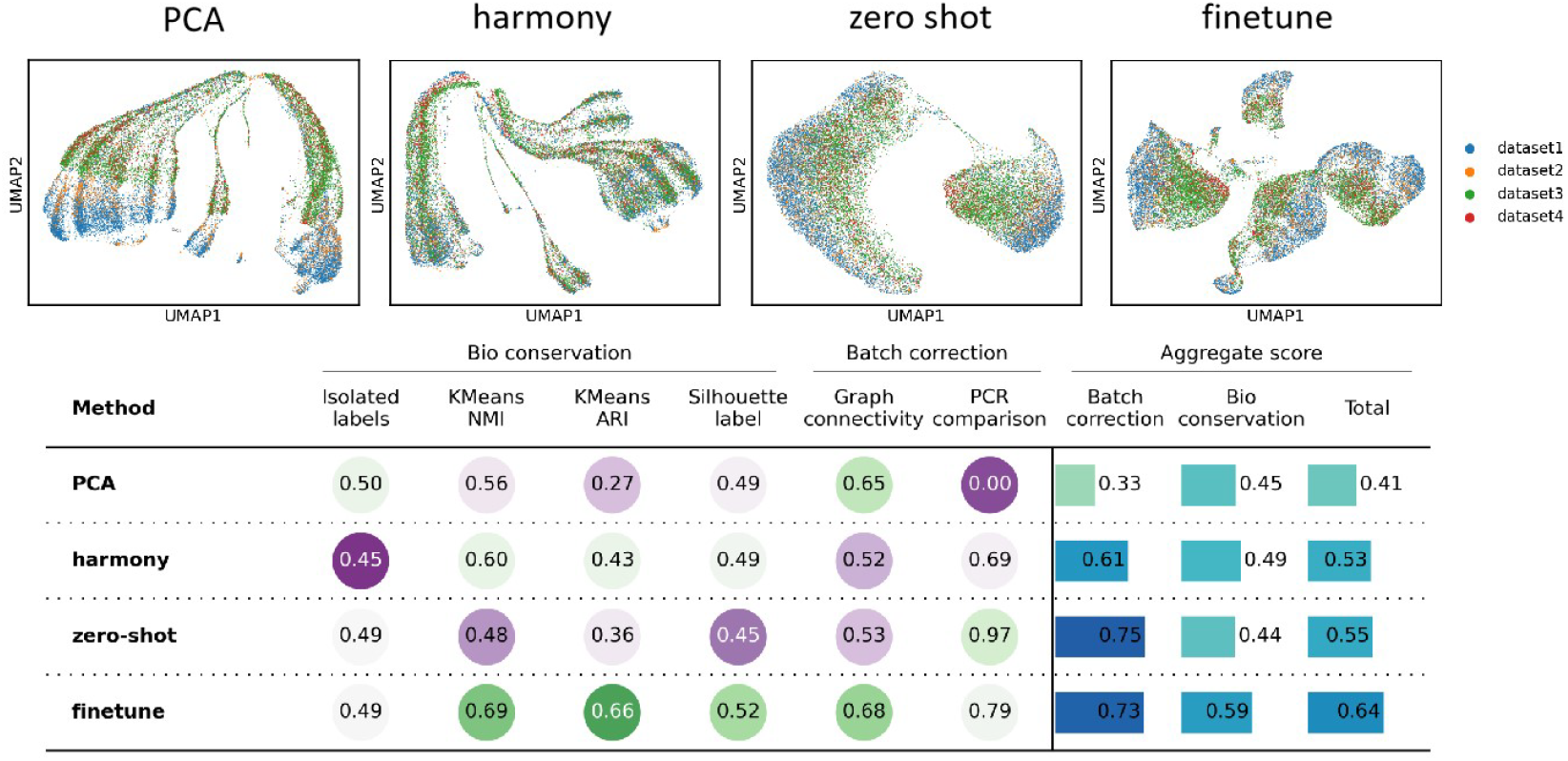
The performance of the CLM-access model in removing batch effects

#### 4.3.2 Experimental Results of Cell Type Annotation

Cell type annotation is a fundamental task in single-cell analysis, playing a crucial role in accurately characterizing the cellular composition and heterogeneity of biological samples. To rigorously assess the performance of CLM-access on cell type annotation, we selected two large-scale datasets—GSE219281 and GSE181346—each comprising approximately 70,000 cells. Traditional tools often struggle with cell type annotation at this scale, as many lack the capacity to efficiently process such large and complex datasets. These two datasets were therefore chosen to provide a challenging and representative benchmark for evaluating CLM-access. We compared CLM-access with a conventional annotation tool scATAnno[34] as well as the state-of-the-art deep learning-based method Cellcano [35]. As illustrated in Table 5, the fine-tuned CLM-access model demonstrates outstanding performance across multiple key evaluation metrics, including accuracy, precision, recall, and micro-averaged F1 score, underscoring its remarkable efficacy and effectiveness in cell type annotation tasks.

**Table 5:** The performance of the CLM-access model in cell type annotation.

| dataset | model | accuracy | precision | recall | macro_f1 |
| --- | --- | --- | --- | --- | --- |
| GSE219281 | CLM-access | **0.7635** | **0.6687** | **0.4795** | **0.5437** |
|  | scATAnno | 0.7283 | 0.5366 | 0.3338 | 0.3610 |
|  | Cellcano | 0.7064 | 0.4685 | 0.3309 | 0.3434 |
| GSE181346 | CLM-access | **0.7470** | **0.8335** | **0.6186** | **0.6742** |
|  | scATAnno | 0.6834 | 0.7129 | 0.5221 | 0.5473 |
|  | Cellcano | 0.6286 | 0.5069 | 0.3871 | 0.3955 |

#### 4.3.3 Experimental Results of RNA Expression Prediction

The prediction of RNA expression from scATAC-seq not only broadens the utility of chromatin accessibility data in transcriptome-level analyses, but also serves as a direct reflection of the model’s capacity to extract and interpret meaningful regulatory signals from peak-level information. The CLM-access model is designed to capture the complex interrelationships among chromatin regions while preserving complete peak-level information within each local patch during data processing. In this study, we use a paired scRNA-seq and scATAC-seq data (see data detail in supplementary file) to examine the performance of CLM-access on RNA expression task, and compare it with two well-known methods Babel [36] and MultiVI[30].

As illustrated in Table 6, Our fine-tuned CLM-access model exhibits great performance in gene expression prediction tasks, with the predicted transcriptomic profiles showing strong Pearson correlation of 0.9175 with the corresponding RNA-seq data. Furthermore, CLM-access consistently outperforms Babel and MultiVI, demonstrating superior performance over current state-of-the-art methods.

**Table 6:** The performance of the CLM-access model in gene expression prediction.

| Model | Pearson | Rmse |
| --- | --- | --- |
| CLM-access | **0.9175** | **1.4185** |
| babel | 0.9101 | 1.5316 |
| MultVI | 0.8845 | 3.5755 |

#### 4.3.4 Experimental Results of Multi-modal Integration

The integration of RNA-seq and ATAC-seq modalities is a critical task in single-cell multi-omics analysis, offering a more comprehensive understanding of cell identities, states, and regulatory mechanisms than single-modality approaches. To evaluate the performance of CLM-access in multimodal integration, we first fine-tuned the model to predict scRNA-seq profiles from scATAC-seq data. The predicted gene expression was then combined with the original RNA-seq data to construct a unified transcriptomic representation, which was subsequently tested on a paired scRNA-seq and scATAC-seq dataset (see details in supplementary file). As illustrated in Figure 3, the result shows integrated features by CLM-access fine-tune method achieve better performance in cell type identification over the snapATAC2 method performed on the scATAC-seq data alone, proving its ability in multi-modal integration.

**Figure 3.**
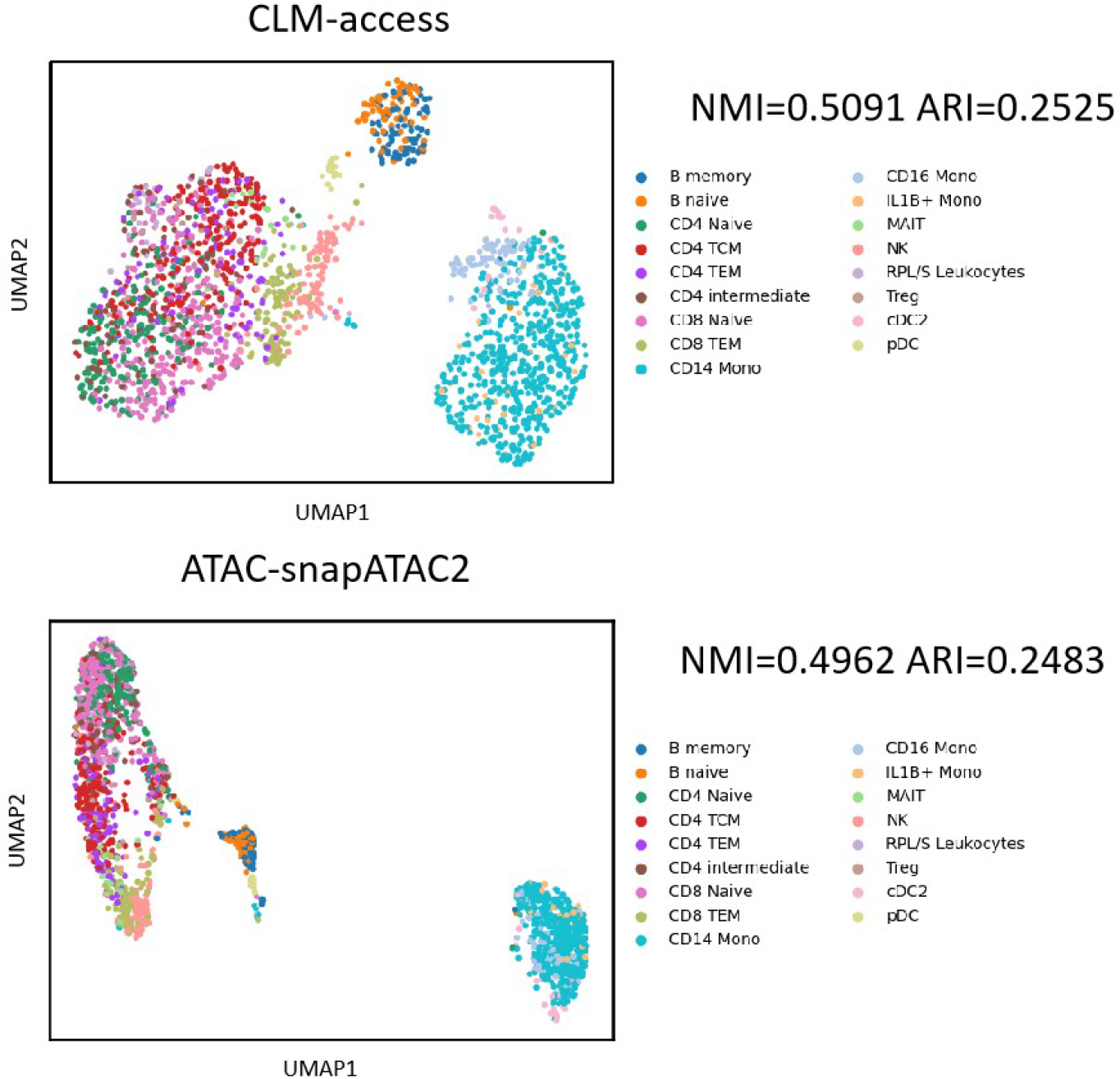
The performance of the CLM-access model in multi-modal integration

## 5 Discussion

We designed CLM-access, a foundation model specifically for scATAC-seq data. Unlike previous foundation models for transcriptomes, CLM-access introduced a novel integration strategy embedded within the Transformer framework: the genomic peaks are partitioned into 2,000 fixed patch regions, with each region treated as an independent token. The encoder comprehensively encodes the peak information within each patch into low-dimensional vectors. After fusing the embeddings of each token with the low-dimensional representations of the peaks as model input, this input is fed into the Transformer for deep learning to reconstruct masked peaks. This approach provides a new perspective for integrating scATAC data into the Transformer architecture. It not only preserves all scATAC peak information without loss but also significantly enhances model performance and training/inference efficiency, overcoming the limitations of existing scATAC-seq-based methods, such as weak framework generalizability, inadequate utilization of large-scale datasets, and the inability to capture all accessible cis-regulatory elements (cCREs).

## Supporting information

Supplement the technical documentation

## Limitations and Future Work

Our current work studied only data of human cells. In our future work, we will further integrate multi-source information to unify cCREs across multiple species within a shared embedding space. Our current model relies solely on chromatin accessibility data, and its attention mechanism covers only regulatory networks at the patch level. We plan to expand the attention coverage to the regulatory network modeling to the cCRE level. To investigate implications and potentials of multi-omics data, we will incorporate more omics modalities, such as RNA-seq, to construct multi-omics “cell sentences,” enabling a more comprehensive characterization of cellular heterogeneity.

