## Supplement the technical documentation for "CLM-access: A Specialized Foundation Model for High-dimensional Single-cell ATAC-seq analysis"

### 1 Source of pre-training data

In this study, we meticulously constructed a dataset named Human-scATAC specifically designed for the pre-training of a foundational model tailored for single-cell chromatin accessibility sequencing (scATAC-seq). This dataset integrates approximately 2.8 million scATAC-seq sequencing reads, with its core components being the CATlas and Descartes databases[1, 2].

Specifically, we employed the chromatin accessibility atlas CATlas, published in 2021, as a pivotal source of pre-training data for the single-cell ATAC (scATAC) analysis framework. The CATlas atlas conducted in-depth analyses on over 1.3 million cell nuclei derived from 222 types of human fetal and adult cell types, successfully identifying around 1.2 million candidate cis-regulatory elements (cCREs). By integrating single-cell chromatin accessibility sequencing (sci-ATAC-seq) data from 30 adult tissues and 15 fetal tissues, this atlas finely dissected cell-type-specific regulatory programs and possessed the capability to systematically interpret non-coding variants associated with complex traits. During the pre-training process, we fully leveraged this resource, aiming to accurately capture the dynamic characteristics of chromatin accessibility across tissues and life stages. The immense scale and high diversity exhibited by the CATlas atlas laid a solid foundation for training models with broad generalization capabilities in the field of single-cell epigenomics.

The Descartes fetal chromatin atlas is also an indispensable and vital component of the Human-scATAC dataset. Leveraging the sci-ATAC-seq3 technology, this atlas performed chromatin accessibility analyses on 790,957 single cells from 15 human fetal organs (ranging from 8 to 20 weeks of gestation). Based on this analysis, a total of 1.05 million regulatory elements, 54 cell types exhibiting transcription factor (TF) motif enrichment characteristics, and cell-type-specific heritabilities corresponding to 34 traits were identified.

Firstly, the Descartes dataset includes fragment data, whereas the CATlas dataset only provides cell-peak matrices. This difference poses an initial challenge for direct comparison and analysis between the datasets. More crucially, the two datasets were processed using different genome reference versions. Specifically, the CATlas dataset was aligned based on the hg38 genome version, while the fragment data in the Descartes dataset was based on the hg19 genome version.

To overcome the issue of inconsistent genome versions, we utilized the liftover toolkit to convert the data in the Descartes dataset from the hg19 coordinate system to the hg38 coordinate system. This step is fundamental for ensuring that the two datasets can be compared and analyzed under the same genomic background[3].

Subsequently, to further align the data format of the Descartes dataset with that of the CATlas dataset, we employed the snapATAC2 toolkit[4]. Using this toolkit, we converted the fragment data in the Descartes dataset into cell-peak matrices, which are consistent with the data format of the CATlas dataset.

Finally, during the data preprocessing stage, we also removed some non-informative candidate cis-regulatory elements (cCREs) to ensure the accuracy and reliability of the dataset. After this series of processing and refinement, the final dataset contains 1,154,464 cCREs, providing a solid foundation for subsequent in-depth analysis and comparison.

### 2 Data used in the pre-training experiment

In this study, we conducted diverse explorations and attempts regarding different pre-training schemes. Given that pre-training using the Human-scATAC dataset would consume substantial computational resources, we introduced the scATAC-seq portion from a pre-compiled pseudo-paired fetal atlas dataset called scCLIP data, which was organized by others earlier[5].

This dataset was constructed to address the scarcity of large-scale, truly paired fetal multi-omics data. It integrates single-cell RNA sequencing (scRNA-seq) and single-cell chromatin accessibility sequencing (scATAC-seq) atlas data derived from fetal organs. To achieve pseudo-pairing, we randomly matched cells from the corresponding atlases based on the same cell type annotations provided in the original studies. The resulting dataset comprises 377,134 pseudo-paired cells, covering 36,601 genes and 1,154,464 chromatin accessibility peaks.

In terms of data processing, to maintain consistency with the original compilation approach, we refrained from performing additional filtering or selection operations on gene and peak features. Instead, we directly utilized all available gene and peak features to conduct cross-modal prediction benchmarking.

### 3 Map ATAC peaks to TADs

To effectively aggregate single-cell ATAC-seq signals within topological associating domains (TADs), we designed and implemented a rigorous computational pipeline[6].

First, we precisely extracted peak coordinate information from the input h5ad file (which contains a cell-by-peak count matrix) and saved it properly in BED format. If the input data utilized the hg19 coordinate system, we employed the liftover tool to convert the peak coordinates to the hg38 reference genome. Simultaneously, we filtered the corresponding h5ad file, retaining only those peaks that were successfully mapped[3]. This ensured the accuracy and consistency of data processing.

Subsequently, we utilized the bedtools intersect tool (with the -loj -wao options and a default overlap ratio of 51% between peaks and TADs, which could be flexibly configured according to actual requirements) to perform an intersection analysis between the peak BED file (either in the original hg38 format or the converted format) and a pre-provided TAD annotation BED file[7]. This step generated a mapping file that precisely associated each peak with its corresponding TAD.

Finally, based on the aforementioned mapping relationships, we integrated the peak counts from the h5ad file, performing signal aggregation at the cellular level within each TAD region. To enhance memory utilization efficiency, we processed the data in a chunked manner. The resulting data, namely the cell-by-TAD signal matrix, was saved in a new h5ad file format. Additionally, to facilitate subsequent data tracing and analysis, intermediate files, including the original peak BED file, the peak-TAD intersection BED file, and the peak-to-TAD mapping table, were also properly preserved.

### 4 Pre-training experiment setup and result supplementation

#### 4.1 Binarization Processing

During the data processing for each patch-segmented region, we first summed up the intensity values of all peak signals within each patch. Subsequently, based on this processed result, we conducted a binary comparison experiment. The specific experimental design is delineated as follows: Two experimental groups were established, one comprising data that had undergone binarization processing, and the other consisting of non - binarized data.

For ATAC data, this study employed two regional segmentation strategies: patch-based segmentation and topological associating domain (TAD)-based segmentation. In the patch-based segmentation experiments, we set up three different patch granularity schemes: 1000, 2000, and 5000. The methodology for TAD segmentation is detailed in Section 3.

In the model training phase, considering that training on the entire Human-scATAC dataset would result in excessively long training times and waste computational resources, we selected a subset of scATAC-seq data from the scCLIP dataset for training[5]. The data was divided into a training set and a test set in a 9:1 ratio. During the training process, we utilized 4 NVIDIA A100 GPUs, each with 80GB of memory, for parallel computation. The model was trained for 15 epochs with a batch size of 8. The model parameters are configured as follows: the number of attention layers is set to 8, the count of attention heads is 8, and the dimension of hidden states is 256.

Analysis revealed that when only the sum of all peak signal intensity values was calculated without binarization, the model encountered significant convergence difficulties during training, struggling to effectively capture the internal feature information of each patch region. In stark contrast, after introducing binarization, the model's learning complexity was substantially reduced. Further investigation indicated that as the granularity of the patch regions decreased, the model was able to retain more complete original information during the feature extraction process, thereby demonstrating superior performance.

##### 4.2 Loss Function Comparison Experiment

In this study, we designed and compared two distinct data processing strategies for patch-segmented regions. Strategy 1 involves summing the intensity values of all peak signals within each patch, followed by binarization. Subsequently, we employed Mean Squared Error (MSE) and Binary Cross-Entropy (BCE) as loss functions for model optimization. Strategy 2 entails directly binarizing all peak signals, then inputting them into the model patch by patch in batches, with BCE serving as the loss function. During the experimental process, we systematically compared the performance of these two strategies.

During the model training phase, considering that training on the entire Human-scATAC dataset would lead to excessively long training times and waste computational resources, we opted to use the scATAC-seq data portion from the scCLIP dataset for training[5]. The data was divided into a training set and a test set in an 9:1 ratio. Specifically, the training set comprised approximately 339420 cells, while the test set contained around 37714 cells. For the training process, we employed four NVIDIA A100 GPUs for parallel computation, each with 80GB of memory. The model was trained for 15 epochs with a batch size of 8. The model parameters are configured as follows: the number of attention layers is set to 8, the count of attention heads is 8, and the dimension of hidden states is 256.

Our preliminary experimental results have confirmed that reducing the granularity of patch regions can effectively enhance model performance. However, this optimization strategy has a notable drawback: the summation operation within patches leads to the loss of a substantial amount of original peak signal information. While blindly increasing the number of patches can alleviate information loss to some extent, it results in a sharp increase in the length of the model's input sequence, significantly escalating the time costs associated with model inference and training. In light of this, our study attempted to adopt Strategy 2, which involves directly binarizing all peak signals and inputting them into the model patch by patch in batches.

As presented in the main text, Strategy 2 attained optimal performance with an input sequence length of merely 2000. This strategy not only substantially improved model performance but also significantly shortened the time required for model inference and training, reflecting an effective balance between computational efficiency and feature preservation capabilities.

##### 4.3 Different Forms of Masking

Through systematic experimental design, this study explored the impact of different masking strategies on model performance and subsequently devised two masking approaches: one involves jointly masking patch tokens and peak signals, while the other only masks peak signals. We conducted comparative experiments to systematically evaluate the differential effects of these two masking strategies on the model's feature learning capabilities.

During the model training phase, given that training on the entire Human-scATAC dataset would entail excessively long training times and a waste of computational resources, we opted to select the scATAC-seq data subset from the scCLIP dataset for training purposes[5]. The data was partitioned into a training set and a test set at an 9:1 ratio, with the training set encompassing approximately 339420 cells and the test set around 37714 cells.

In the training process, we leveraged 4 NVIDIA A100 GPUs, each with 80GB of memory, for parallel computing. The model underwent 15 epochs of training, with a batch size of 8. As for the model parameter configuration, we set the number of attention layers to 8, the number of attention heads to 8, and the dimension of hidden states to 256.

As demonstrated in the main text, the masking strategy that simultaneously masks both patch tokens and peak signals substantially elevates the model's learning complexity, rendering it difficult for the model to effectively extract crucial biological features from highly masked input sequences. Based on this result, subsequent experiments adopted the strategy of only masking peak signals. This approach maintained the model's convergence stability while substantially reducing the risk of information loss, ensuring that the model could focus on learning the local features and global distribution patterns of peak signals. This optimized strategy provided crucial support for enhancing the model's performance in subsequent experiments.

##### 4.4 The CLM-access framework demonstrates remarkable zero-shot capabilities

After the pre-training phase, we randomly selected approximately 37,000 data points from the scCLIP dataset as a test set to evaluate the model's zero-shot learning capability. Specifically, we first trained three pre-trained models with different parameter sizes using the entire 2.8 million ATAC data points. Subsequently, to investigate the impact of pre-training data volume on model performance, we further tested the models' performance using 300,000, 1.4 million, and 2.8 million data points, respectively.

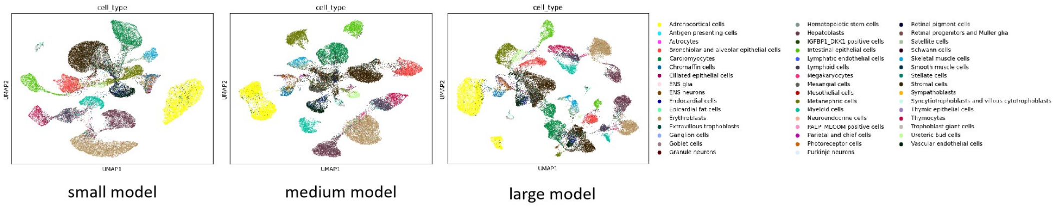


Figure 1: The zero-shot clustering performance of the model under different parameters

###### 4.4.1 The performance under different parameter sizes.

Given that the parameter scale of the Transformer architecture significantly impacts model performance in large language models, this study conducted a systematic experimental design to compare the model performance under three distinct parameter configurations. Specifically, we set the number of Transformer layers, the quantity of multi-head attention heads, and the dimension of the intermediate layer to [8, 8, 256], [8, 8, 512], and [12, 8, 512], respectively.

It is worth noting that for all three parameter settings, we utilized the complete Human-scATAC dataset. Consequently, during the training process, we employed 4 NVIDIA A100 GPUs, each with 80GB of memory, for parallel computation. All models were trained for 15 epochs with a batch size of 32.

As illustrated in Figure Figure 1 , the experimental results indicate that the model with the parameter configuration of [8, 8, 512] performed optimally in the performance evaluation. Further increasing the parameter scale did not lead to performance improvements but instead exhibited a trend of diminishing marginal returns.

Based on the aforementioned observations, this study hypothesizes that this phenomenon may be related to the scale and distributional characteristics of the pre-training dataset. Under conditions of limited data volume, excessively expanding model capacity may increase the risk of overfitting, thereby constraining generalization ability. Therefore, taking into account both computational efficiency and model performance, this study ultimately selected [8, 8, 512] as the standard parameter configuration for the Transformer component. This strategy effectively controls computational resource consumption while maintaining the model's learning capabilities.

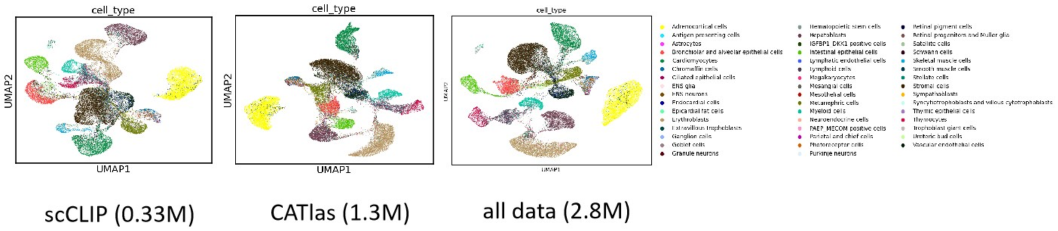


Figure 2: The zero-shot clustering performance of the model under different data size

###### 4.4.2 The performance under different data volumes.

In our prior research, we underscored the potential impact of pre-training data scale on model performance. Building upon this hypothesis, the present study adopted a controlled variable approach to design the experiments.

Specifically, we fixed the number of Transformer layers, the count of multi-head attention heads, and the dimension of the intermediate layer at [8, 8, 256]. Notably, while maintaining the same parameter quantity, we employed three distinct datasets: the unutilized training subset of scATAC-seq data from the scCLIP dataset, the CATlas dataset, and our curated comprehensive Human-scATAC dataset. These datasets comprised approximately 330,000, 1.4 million, and 2.8 million samples, respectively.

During the training process, we harnessed the computational power of 4 NVIDIA A100 GPUs, each equipped with 80GB of memory, for parallel computing. All models underwent 15 epochs of training with a batch size of 32.

As illustrated in the Figure 2, the experimental results unequivocally demonstrate that, under the premise of a consistent model architecture, as the volume of pre-training data progressively expands, the model's performance metrics on downstream tasks exhibit a discernible upward trajectory. Moreover, this improvement is approximately linearly proportional to the data scale. This finding substantiates the notion that large-scale data enhances the feature representation capability of deep learning models, thereby offering a theoretical foundation for dataset construction strategies in subsequent experiments.

### 5 Experimental setup and result supplementation for downstream tasks

The pre-trained CLM-access model generates high-quality cellular representations by modeling ATAC (Assay for Transposase-Accessible Chromatin) sequencing data. As illustrated in Figure 1(c), this model possesses remarkable adaptability for various downstream applications in single-cell analysis through supervised fine-tuning. Specifically, it can be applied to tasks such as batch correlation analysis, cell type annotation and gene expression prediction.

#### 5.1 Downstream analysis model structure

The pre-trained CLM-access model generates high-quality cellular representations by modeling data from Assay for Transposase-Accessible Chromatin (ATAC) sequencing. Following supervised fine-tuning, this model demonstrates exceptional adaptability across a diverse range of downstream tasks in single-cell analysis, encompassing batch effect correction, cell type annotation, gene expression prediction, and multi-modal integration.

In the batch effect removal task, when conducting peak expression prediction, we introduce a batch encoder. The output features from this batch encoder are concatenated with those generated by the Transformer. Subsequently, the concatenated features are fed into a unified decoder, while other settings (including the loss function) remain consistent with those used during the pre-training phase.

For the cell type annotation task, we fine-tune the model using a reference dataset equipped with ground truth labels. The core objective of fine-tuning is to minimize the classification loss for cell type prediction. By optimizing this objective, we enhance the model's capability to accurately assign cell types based on the input features.

In the gene expression prediction task, we fine-tune the model using a paired dataset that integrates both RNA-seq and ATAC-seq modalities. To rigorously evaluate its predictive performance, we validate the model on a held-out test set, adhering to established benchmarking protocols. During the fine-tuning phase, we incorporate a gene expression decoder module, which is specifically designed to output gene expression profiles. To prevent the propagation of biases that may arise during the pre-training process, this module is randomly initialized. Its architecture comprises a fully connected neural network that takes as input the feature representation extracted from the [CLS] token (a contextualized sequence embedding) and outputs predictions for 2,000 pre-selected highly variable genes from the dataset. The primary goal of fine-tuning is to minimize the predictive discrepancy between the model's gene expression estimates and the ground truth measurements. This discrepancy is quantified through a task-specific loss function. By optimizing this loss metric, we aim to systematically reduce the prediction error, thereby improving the model's ability to infer accurate gene expression patterns directly from the input features.

In the multi-modal integration task, we fine-tune the model using an approach analogous to the gene expression prediction task. Specifically, we sum the predicted RNA sequences with the ground-truth RNA sequences, thereby achieving information integration across the ATAC-seq and RNA-seq modalities. This method effectively facilitates the fusion of information from both modalities.

#### 5.2 Batch Correction

Batch effects refer to the observed discrepancies in gene or peak expression data, which arise from technical variations between different batches of samples processed at distinct time points or under varying laboratory conditions. These variations can potentially obscure the genuine biological differences among single cells. To address this issue, CLM-access constructs feature vectors for each batch and conducts fine-tuning, thereby aiding in the elimination of batch effects.

In our proposed methodology, we leverage all the weights from the pre-trained model to initialize the fine-tuning model. This ensures that the fine-tuning model benefits from the knowledge acquired during the pre-training phase. For peak expression prediction, we incorporate a batch encoder, concatenating its output features with those generated by the Transformer. The combined features are then fed into a unified decoder, while other settings, including the loss function, remain consistent with those used during pre-training. This approach is designed to better preserve the knowledge learned during pre-training.

As demonstrated in the main text, we conducted a comparative evaluation of our model against other benchmark methods, including Principal Component Analysis (PCA) and Harmony. The results clearly indicate that our model outperformed these traditional approaches across all performance metrics. PCA (Principal Component Analysis) is a dimensionality reduction technique that integrates samples without implementing any batch correction procedures. Consequently, PCA outputs inherently preserve the original batch-specific variations present in the data. For this study, we derived the PCA embeddings of cells using the scIB.integration.harmony function from the Python package scIB (version 1.1.7) [8].

Harmony, in contrast, is a single-cell batch correction method that employs iterative soft clustering to align cells across diverse batches[9]. It operates by initially projecting cells into a shared low-dimensional space via PCA, followed by iterative refinement of cell embeddings to minimize batch effects while safeguarding biological heterogeneity. To compute the Harmony latent space, we utilized the scib.integration.harmony function within the scIB Python package (version 1.1.7) [8], utilizing the raw count matrix as input. This strategy facilitated effective batch harmonization, enabling a more accurate and biologically relevant analysis of the data.

Regarding parameter settings, we adopt the pre-trained parameters with the Transformer architecture configured as [8 layers, 8 attention heads, and a hidden state dimension of 256], trained on the entire Human-scATAC dataset. Our empirical findings indicate that increasing the parameter scale does not lead to a significant improvement in clustering performance. Therefore, to optimize the fine-tuning time and efficiency, we opted for the pre-trained model with the aforementioned parameter configuration. During the fine-tuning process, we adjust all parameters and utilize a single NVIDIA A40 GPU with 48GB of memory. The model undergoes 300 epochs of training, with a batch size of 8.

The test data employed in this study are derived from the single-cell multi-omics Benchmark [10], comprising four distinct subsets generated using the 10x Multiome sequencing technology to ensure consistency and comparability across datasets. The first subset, Dataset 34, contains 10000 peripheral blood mononuclear cells (PBMCs) with granulocytes removed, obtained through cell sorting.The second subset, Dataset 35, consists of 3000 PBMCs, also depleted of granulocytes via cell sorting. The third subset, Dataset 36, encompasses 10000 PBMCs without any cell sorting, thus preserving the natural cellular heterogeneity. The fourth subset, Dataset 37, features 3000 PBMCs, again without cell sorting. These four datasets were integrated to form a comprehensive test dataset, offering a diverse range of cellular compositions and sequencing depths. The integration process involved standardizing the data preprocessing pipelines to ensure uniformity in feature extraction and normalization, thereby enabling a robust evaluation of the model's performance across varying cellular contexts and data scales.

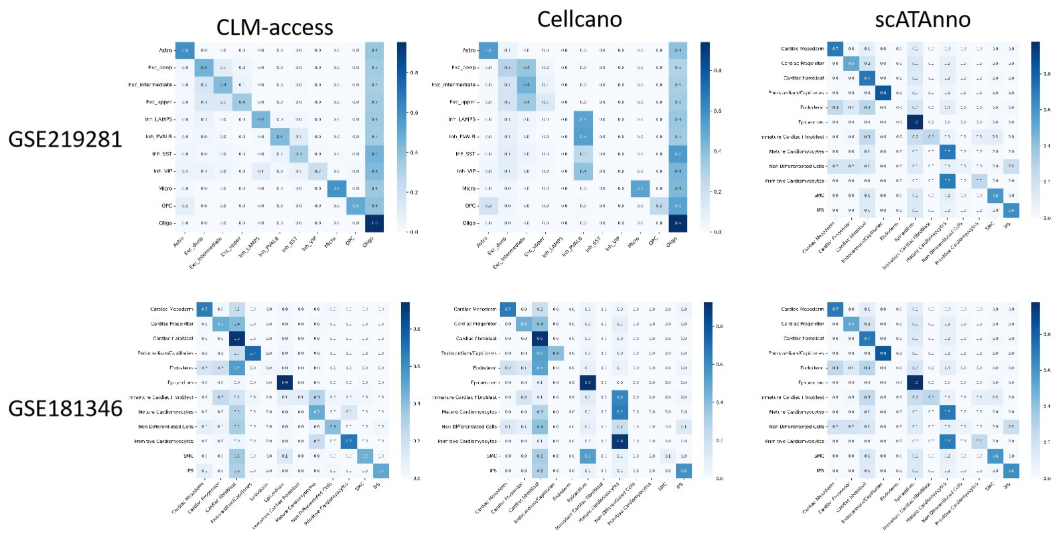


Figure 3: Fine-tune the performance of the CLM-access model in cell type annotation

##### 5.3 Cell Type Annotation

For the cell type annotation task, we fine-tuned our model using datasets GSE219281 and GSE181346, which are equipped with ground-truth labels. To evaluate the annotation performance, we allocated 70% of each dataset for training and reserved the remaining 30% for testing.

During the fine-tuning process, we initialized the model using all the weights from the pre-trained model. However, the output cell type classifier was randomly initialized to facilitate learning cell types specific to our datasets.

The primary objective of fine-tuning was to minimize the classification loss in cell type prediction. By optimizing this goal, we aimed to enhance the model's capability to accurately assign cell types based on input features.

In summary, this approach enabled us to effectively leverage the pre-trained model while adapting the output layer to the specific cell types present in our reference datasets, ultimately improving annotation performance on the query datasets.

Regarding parameter settings, we employed the pre-trained parameters with the Transformer architecture configured as [8 layers, 8 attention heads, and a hidden state dimension of 256], trained on the complete Human-scATAC dataset. Our empirical analysis revealed that increasing the parameter scale did not lead to a substantial improvement in clustering performance. Therefore, to optimize the fine-tuning time and efficiency, we opted for the pre-trained model with the aforementioned parameter configuration. Throughout the training process, we fine-tuned all parameters using a single NVIDIA A40 GPU with 48GB of memory. The model was trained for 20 epochs, with a batch size of 8.

As illustrated in Figure 3, our fine-tuned model, CLM-access, outperformed the conventional scATAC-seq cell type annotation model, Cellcano, across both datasets. The gene score matrix for Cellcano was generated using snapATAC2, with all other parameters set to their default values.

##### 5.4 Gene Expression Prediction

In the gene expression prediction task, we fine-tuned the model using a paired dataset that integrates two modalities: RNA sequencing (RNA-seq) and transposase-accessible chromatin sequencing (ATAC-seq). To rigorously evaluate its predictive performance, we adhered to established benchmarking protocols and validated the model on a fixed test set.

During the fine-tuning phase, we initialized the model weights using all parameters from the pre-trained model, with one critical exception: the gene expression decoder module. This decoder was specifically designed to output gene expression profiles. To prevent bias propagation from the

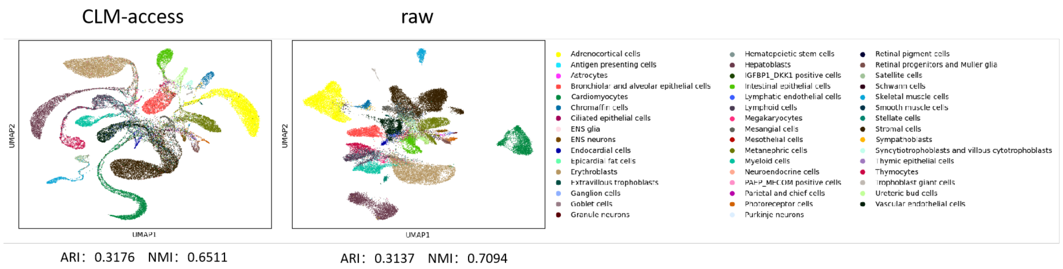


Figure 4: The performance of gene expression prediction on scCLIP data

pre-training process, we randomly initialized it. Its architecture comprises a fully connected neural network that takes feature representations extracted from the [CLS] token (contextualized sequence embeddings) as input and outputs predictions for specified genes. Notably, the number of target genes can be configured as a hyperparameter, enabling task-specific adaptability. The fully connected layer structure of the decoder facilitates learning cross-modal mappings between RNA-seq and ATAC-seq features, thereby capturing the regulatory synergies embedded in the paired omics data.

The primary objective of fine-tuning was to minimize the discrepancy between the model's predicted gene expression values and the ground-truth measurements through a task-specific loss function. By optimizing this loss metric via gradient-based updates, we systematically enhanced the model's ability to infer accurate gene expression patterns directly from input features. This optimization process inherently improved the model's interpretability, enabling it to emphasize biologically meaningful regulatory relationships while excluding technical confounders.

Regarding parameter settings, we adopted pre-trained parameters with a Transformer architecture configured as [8 layers, 8 attention heads, and a hidden state dimension of 256], trained on the complete Human Single-Cell Chromatin Accessibility Sequencing (Human-scATAC) dataset. Our empirical findings revealed that increasing the parameter scale did not significantly improve clustering performance. Therefore, to optimize fine-tuning time and efficiency, we utilized the pre-trained model with the aforementioned parameter configuration. During training, we fine-tuned all parameters using an NVIDIA A40 GPU with 48GB of memory. The model was trained for 150 epochs with a batch size of 8.

As demonstrated in the main text, the data employed for both training and testing were derived from the first batch of the merged batch data mentioned in Section 5.2, with 80% allocated for training and 20% for testing. We conducted a comparative analysis between our model and other modality translation models. For these comparisons, predictions were performed using full-length genes, and the data source remained consistent, utilizing the first batch of the merged batch data specified in Section 5.2.

#### 6 The indicators used in this article

In this study, we employed a comprehensive suite of evaluation metrics to thoroughly assess model performance, clustering quality, and the structural characteristics of the data. Below is a concise summary of these metrics:

Accuracy: Accuracy denotes the proportion of samples correctly predicted by the model relative to the total number of samples. This metric serves as a fundamental gauge for evaluating the overall predictive accuracy of classification models.

Precision: Precision quantifies the proportion of samples actually belonging to the positive class among all samples predicted as positive. It focuses on the accuracy of positive predictions, particularly crucial in scenarios where minimizing false positives is essential.

Recall: Recall measures the proportion of positive samples correctly identified by the model out of all actual positive samples. This metric assesses the model's ability to detect all positive instances, which is of paramount importance in tasks requiring exhaustive identification of positive cases, such as disease diagnosis.

Macro_F1: Macro_F1 represents an arithmetic mean of F1 scores across all classes, where the F1 score is the harmonic mean of precision and recall. By integrating both precision and recall, Macro_F1 offers a holistic evaluation of model performance on imbalanced datasets, providing a more nuanced reflection of performance across various classes.

Isolated Labels: Isolated labels typically refer to labels or samples in clustering or classification tasks that exhibit weak associations with other classes or clusters. These labels or samples may represent outliers or noise, and their identification and handling contribute to enhancing the model's robustness.

KMeans NMI (Normalized Mutual Information): NMI quantifies the similarity between two clustering results by assessing the consistency between KMeans clustering outcomes and ground-truth labels. A higher NMI value indicates greater similarity between the clustering results and the true labels, thereby providing a quantitative measure of clustering algorithm performance.

KMeans ARI (Adjusted Rand Index): ARI is another metric for evaluating the similarity between two clustering results, adjusted for randomness. It offers a more precise assessment of KMeans clustering quality, with a higher ARI value signifying a closer alignment between the clustering results and the actual data structure.

Silhouette Label: The silhouette coefficient evaluates the cohesion and separation of individual samples within clusters. By calculating the silhouette coefficient for each sample, it enables a quantitative assessment of clustering quality, where a higher coefficient denotes superior clustering efficacy.

PCR comparison involves a systematic quantitative evaluation of core parameters, including cycle threshold (Ct) values, amplification efficiency (E), sensitivity, and detection consistency, to assess differences in amplification performance, quantitative accuracy, and biological relevance across various PCR experimental conditions (e.g., primer/probe design optimization, reaction system refinement), technical methodologies (e.g., qPCR vs. digital PCR), or predictive models . This approach provides critical data support for experimental design optimization, methodological validation, and model reliability assessment.

Graph Connectivity: Graph connectivity measures the degree of node interconnection within a graph, serving as a pivotal metric for assessing graph structural properties. In graph clustering or analysis, evaluating graph connectivity aids in understanding cluster or community structures, thereby facilitating a deeper comprehension of the inherent relationships within the data.

These metrics collectively offer a multifaceted evaluation of model performance, clustering quality, and data structural characteristics, furnishing robust support for in-depth analysis and model optimization.

[5] Lei Xiong, Tianlong Chen, and Manolis Kellis. scclip: Multi-modal single-cell contrastive learning integration pre-training. In *NeurIPS 2023 AI for Science Workshop*, 2023.
